# Efficient randomization of biological networks while preserving functional characterization of individual nodes

**DOI:** 10.1101/069245

**Authors:** Francesco iorio, Marti Bernardo-Faura, Andrea Gobbi, Thomas Cokelae, Giuseppe Jurman, Julio Saez-Rodrigue

**Author notes:** Equal contributors (Francesco iorio, Marti Bernardo-Faura, Andrea Gobbi, Thomas Cokelae).

## Abstract

**Background:** Networks are popular and powerful tools to describe and model biological processes. Many computational methods have been developed to infer biological networks from literature, high-throughput experiments, and combinations of both. Additionally, a wide range of tools has been developed to map experimental data onto reference biological networks, in order to extract meaningful modules. Many of these methods assess results’ significance against null distributions of randomized networks. However, these standard unconstrained randomizations do not preserve the functional characterization of the nodes in the reference networks (i.e. their degrees and connection signs), hence including potential biases in the assessment.

**Results:** Building on our previous work about rewiring bipartite networks, we propose a method for rewiring any type of unweighted networks. In particular we formally demonstrate that the problem of rewiring a signed and directed network preserving its functional connectivity (F-rewiring) reduces to the problem of rewiring two induced bipartite networks. Additionally, we reformulate the lower bound to the iterations’ number of the switching-algorithm to make it suitable for the F-rewiring of networks of any size. Finally, we present *BiRewire 3*, an open-source Bioconductor software enabling the F-rewiring of any type of unweighted network. We illustrate its application to a case study about the identification of modules from gene expression data mapped on protein interaction networks, and a second one focused on building logic models from more complex signed-directed reference signaling networks and phosphoproteomic data.

**Conclusions:** *BiRewire3* it is freely available at https://www.bioconductor.org/packages/BiRewire/, and it should have a broad application as it allows an efficient and analytically derived statistical assessment of results from any network biology tool.

## 1 Background

Representing and modeling biological processes as networks, in particular signaling and gene regulatory relations, is a widely used practice in bioinformatics and computational biology. This bridges these research fields to the vast repertoire of tools and formalisms provided by graph-and complex-network-theory. Furthermore, this facilitate an integrative analysis of experimental observations, either by derivation of networks from the data, or by mapping the latter on the former. Hence, network-based approaches have become a popular paradigm in computational biology [1, 2].

In the last few years this has allowed the design of a broad assortment of algorithms and tools whose aim ranges from providing an interpretative framework for the modeled biological relations, to the identification of network-modules able *to explain* phenotypic traits and experimental data from large reference signaling graphs [3, 4]. Many methods in this last class aim at identifying a sub-network, for example, that is composed by the most differentially expressed or significantly mutated genes [5, 6, 7, 8, 9], or that it is targeted by a given external perturbation [10, 11, 12, 13, 14]. Toward this aim different optimization procedures have been used to analyze experimental data, identifying a pathway that is deregulated in a given disease, or whose activity is perturbed upon a given drug treatment.

In many approaches, directed signed networks (DSNs, formally defined in the following sections) are used to model pathways and to interlink pathways from a given collection. In these networks, nodes represent biological entities (typically proteins) while edges represent the biological relationships between them (e.g., the activity of protein A affects that of protein B). These edges have a direction to discriminate effectors and affected nodes in a modeled relation, and a sign to specify whether the modeled relation is an activation (positive sign) or an inhibition (negative sign).

Unsigned/undirected edges modeling generic interactions can be also present. When available, sign and direction allow a more detailed detection of the nature of the interaction between the nodes. In this study, the number, sign and direction of a node’s connections are cumulatively denoted by the *functional characterization level* (FCL) of the corresponding modeled biological entity (from now entity).

In a reference network modeling a set of interlinked pathways or protein-protein-interactions, the FCL might be high for a node that models a *functional hub.* For example a kinase phosphorylating a large number of substrate proteins will have a high number of outgoing edges with positive sign. Similarly, a gene activated by a large number of transcription factors will have a high number of positive in-coming edges. On the other hand the FCL might be strongly biased by the relevance of a biological entity in a given research field, and the resource the network has been assembled from. For example, in a cancer focused reference network it is reasonable to find nodes that have a high FCL just because they have oncogenetic or tumor-suppressive properties, thus have been studied more than others. As a consequence, solutions to the network optimization problems tackled in bioinformatics (and mentioned above) can be strongly influenced by the topology of the initial network, and by the FCL of its nodes.

In an attempt to overcome this issue, some tools assess this bias by comparing their provided sub-network solutions with those that would be obtained (using the same experimental data and the same algorithm) across a large number of trials, each starting from a reference network that is a randomized version of the original one. Many other tools neglect this aspect and the significance of the solution is computed by randomizing the experimental data only. For both options, the expectation of some topological properties (for example the inclusion of a given edge or node) of the sub-network solutions is estimated by analyzing the random solutions obtained across the trials. In this way, the significance of these properties is quantified as the divergence from their expectation, testing against the null hypothesis that there is no association between the analyzed experimental data and the outputted sub-network solutions.

To our knowledge, all the existing methods assessing their solution significance through reference network randomizations make use of a simple edge shuffling. This means that in a randomization trial each edge of the network is simply set to link two randomly selected nodes. This implicitly means that null models resulting from this randomization strategy are totally unconstrained with regards to the degree of the nodes, and the way they are linked to each other in the original network. Therefore, the impact of the FCL of the nodes in the original reference network on the outputted sub-network solution is not considered. In order to take this into account a constrained randomization strategy preserving the FCL of all the nodes in the original network must be used.

The problem of randomizing an undirected and unweighted network while preserving the degree of its nodes, i.e. the total number of incident edges for each node, is known in graph theory as *network rewiring* and unfortunately presents itself with analytical and numerical challenges [15]. With the additional constrain that the network to rewire is bipartite (i.e. nodes can be partitioned into two sub-sets such that there are no edges linking nodes in the same set), this problem reduces to randomizing a binary matrix preserving its marginal totals, i.e. its row-wise and column-wise sums. Several algorithms exist to solve this problem [16, 17] but the computationally efficient randomization of moderately large matrices (therefore the rewiring of large bipartite networks) is still challenging. Additionally, to our knowledge, none of the methods published is formally shown to be able to actually simulate samplings from the uniform distributions of all the possible binary matrices with prescribed marginal totals. Such proof exists for methods rewiring directed binary networks based on *swap-and-fill* strategies applied to their adjacency matrices [18] but not dealing with DSNs. Finally, some recent methods have been proposed to solve the related (but yet different from FCL preserving rewiring) problem of randomizing metabolic networks in a mass-balanced way [19].

In [20] we showed how an algorithm based on a Monte Carlo procedure known as the *switching-algorithm* (SA) [21] can be used to efficiently randomize large cancer genomics datasets preserving the mutation burdens observed across patients and the number of mutations harbored by individual genes (hence to efficiently rewire large bipartite networks). To this aim, we derived a novel lower bound for the number of steps required by the SA in order for its underlying Markov chain to reach a stationary distribution. Additionally, we implemented the SA in the R package *BiRewire* (publicly available on Bioconductor [20]) and we showed a massive reduction in computational time requirements of our package and bound with respect to other existing R implementations [22] and bounds [21].

Here (i) we introduce the problem of rewiring a DSN modeling a biological network in a way that the FCL of all the modeled entities is preserved: *F-Rewiring;* (ii) we formally show how this problem reduces to rewiring 2 bipartite networks; (iii) we provide a generalized bound to the SA for bipartite networks of any size; and (iv) we show the validity of the Markov chain convergence criteria (used in our previous work) for F-rewiring DSNs.

Finally, we provide an overview of the functions included in a new version of *BiRewire* for F-Rewiring, and we show results from two case studies where solutions obtained with two network optimization methods (BioNet [9], and CellNOpt [23]) are assessed for statistical significance and intial reference network biases against constrained null models generated with *BiRewire*.

## 2 Methods

### 2.1 Preliminary notations

The problem we are tackling is the computationally efficient randomization of a directed and signed network (DSN) (formally defined below) in a way that some local features of its individual nodes are preserved.

In such a network 𝒢 = (*V, E*), the edges in *E* can be encoded as triplets (a, b, *) where a is called source node, b is called target node and * is a label denoting the sign of the relation occurring among them, which could be positive, * = +, or negative, * = −.

According to this definition, in a DSN the edge (*a, b*, +) is different from the edge (*a, b*, −), thus making this formalism more flexible than that provided by a directed weighted network (with weights ( {+1, −1}). In fact, differently from such a model, in a DSN two edges with same terminal nodes and direction but different sign can coexist. In addition, a DSN is different and less general than a multidigraph (a directed multigraph), because only two possible edges with the same direction can coexist between the same couple of nodes.

Given an edge *e* ( *E*, we define the function λ(*e*): *E* → {+, −}, mapping each edge to its sign label. Given a node v ( *V*, we define its *in-bound-star I*(*v*) as the set of edges in *E* having v as destination, *I* (*v*) = {*e* ∈ *E*: *e* = (*a, v*, *)}. Similarly, considering the edges having *v* as source defines its *outbound-star, O*(*v*) = {*e* ∈ *E*: *e* = (*v,b*, *)}. Imposing as additional condition for an edge to be included in these sets that of having a fixed sign label, defines positive and negative *in-bound* and *outbound stars.* Formally, the *v positive-* (respectively *negative) in-bound-star* is the set of edges in *G* having v as destination and positive (respectively negative) label, *I*^+^(*v*) = {*e* ∈ *I*(*v*): λ(*e*) = +} (respectively *I*^−^(*v*) = {*e* ∈ *I*(*v*): λ(*e*) = −}). Analogously, the *v positive-* (respectively *negative*) *-out-bound-star* is the set of edges in *G* having v as source and positive (respectively negative) label, *O*^+^(*v*) = {*e* ∈ *O*(*v*): λ(*e*) = +} (respectively *O*^−^(*v*) = {*e* ∈ *O*(*v*): λ(*e*) = −}).

By naturally extending the definition of *node degree* (i.e. the number of edges connected to a node) to these formalisms, we call positive-in-degree of a node v the quantity |*I* ^+^(*v*)| equal to the number of edges with positive label having v as destination. Similarly we define the v *negative-in-degree, positive-out-degree* and *positive-in-degree*, the quantities |*I*^−^(*v*)|, |*O*^+^(*v*)| and |*O*^−^(*v*)|, respectively.

In the light of the introduced notation, the object of this study can be redefined as the randomization of the edges of a DNS G while preserving not only its general node-degrees (*network rewiring*), but also all the signed degrees defined above, for all the nodes: *network F-rewiring.*

A biological pathway can be naturally represented through a DNS *𝒢* = (*V, E*). In this case the nodes in *V* would represent biological entities, and the edges in *E* would represent functional relationships occurring among them, whose type would be defined by the sign label (+ for *activatory* and — for *inhibitory* interactions), with directions indicating effector/affected roles (source/destination of the edges). In this case the signed degrees introduced above would define the functional characterization level (FCL) of the individual biological entities considering all the possible roles that they can assume within a given pathway.

Particularly the positive-out-degree of a node *v* would correspond to the level of characterization of the corresponding biological entity as *activator* of other entities; the negative-out-degree would correspond to its characterization as *inhibitor*; finally, the positive-, respectively negative-, in-degree of a node would correspond to the level of characterization of the corresponding entity as *activated*, respectively *inhibited*, by other entities in the same DSN.

As a consequence, the ultimate goal of this study is to efficiently randomize a pathway (or a collection of interlinked pathways) in a way the functional characterization levels of its individual entities, i.e. the signed-directed degrees of all the nodes, are preserved.

### 2.2 F-rewiring of a directed signed networks is reducible to the rewiring of two bipartite networks: reduction proof

Let us consider a directed signed network (DSN) *𝒢* = (*V, E*), with λ(*e*) ∈ *𝒢* {−, +}, ∀*e* ∈ *E* and a transforming function *f* (*𝒢*), from the set of all the possible DSNs to the set of all the possible pairs of bipartite networks (*B*_+_, *B*_−_), such as *B*_*_ = (S_*_, *D*_*_, *E*_*_), whose node sets are defined as *S*_*_ = {*v* ∈ *V*: ∃ (*v*,*x*, *) ∈ *E*}, and *D*_*_ = {*v* ∈ *V*: ∃ (*x, v*, *) ∈ *E*}, with * ∈ {+, −}. Worthy of note is that the same node of *𝒢* can be both a source (therefore belonging the set *S*_*_) for some edge in E, and a destination (therefore belonging to the set *D*_*_) for some other edge in E. As a consequence f should also relabel the nodes (for example adding a subscript to labels of the nodes in *D*_*_). Here, for simplicity we will neglect this relabeling.

As a conclusion, the function *f* maps *𝒢* to two bipartite networks (BNs) (*B*_+_,*B*_−_) such that *B*_+_ = (*S*_+_, *D*_+_, *E*_+_) is the BN induced by the positive edges of *𝒢*, where all the sources of these edges are included in the first node set *S*_+_, all the destinations in the second set *D*_+_ and two nodes across these two sets are connected by an undirected edge if they are connected in the original network *𝒢* by a positive edge that goes from the node in the first set to that in the second one. The second bipartite network of the pair *B*_−_ is similarly induced by the negative edges of *𝒢*. Formally *E*_*_ = {(*s*,*d*): *s* ∈ *S*_*_, *d* ∈ *D*_*_ and ∃ (*s*,*d*, *) ∈ *E*}, with * ∈ {+, −}. An example of this transformation is shown in Figure 1A.

**Figure 1.**
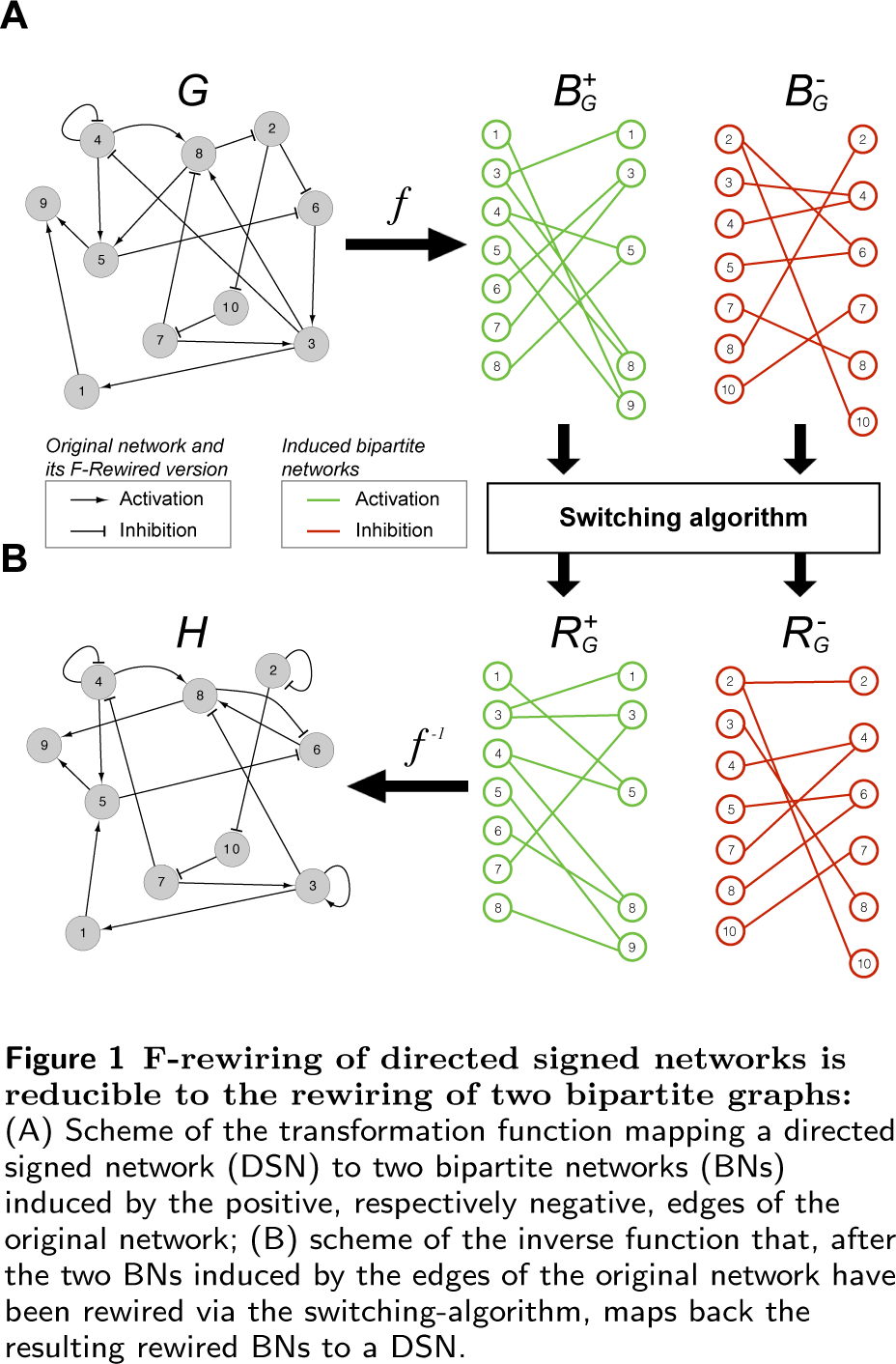
F-rewiring of directed signed networks is reducible to the rewiring of two bipartite graphs: (A) Scheme of the transformation function mapping a directed signed network (DSN) to two bipartite networks (BNs) induced by the positive, respectively negative, edges of the original network; (B) scheme of the inverse function that, after the two BNs induced by the edges of the original network have been rewired via the switching-algorithm, maps back the resulting rewired BNs to a DSN.

It can be shown that such a function *f* realizes a bi-jection between the set of all the possible DNSs and the set of all the possible pairs of BNs [24]. As a consequence its inverse *f*^−1^ is a function from the set of all the possible pairs of BNs to the set of all the possible DSNs, and it is defined as *f* ^−1^(*B*_1_,*B*_2_) = *𝒢* = (*V, E*), where

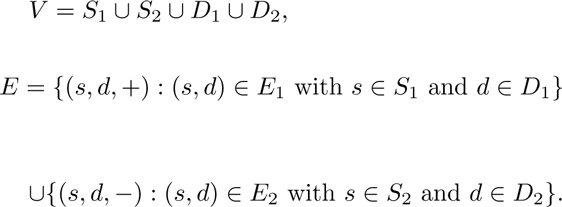

For simplicity, we assume that *f*^−1^ re-assignes to the nodes their original labels before constructing the node/edge sets of *𝒢*, if they were relabeled by the function f. An example of this inverse transformation is shown in Figure 1B.

#### Proposition 2.1.

*Let be 𝒢* = (*V, E*) *a DSN modeling a pathway (or a set of interlinked pathways) P, and f the transformation function described above f*(*𝒢*) = (*B*_+_,*B*_-_). *If R*_+_ *and R*_−_ *are rewired versions of B*_+_ *and B*_-_ *respectively, then f*^−1^(*R*_+_, *R*_−_) = *𝓗* is a *randomized version of 𝒢 in which the signed-directed degrees of all the nodes v* ∈ *V, i.e. the quantities* |*I*^+^(*v*)|, |*I*^−^(*v*)|, |*O*^+^(*v*)|, |*O*^−^(*v*)|, *are kept equal to their original values. This implies that 𝓗 is an F-rewired version of 𝒢, hence a randomization of P in which the functional characterizations of the individual entities are preserved.*

**Proof**. First of all we need to show that *𝓗* is a randomized version of *𝒢*, in other words that *𝓗* is a directed signed network with the same nodes set and number of edges of *𝒢* and the same signed-directed node degrees but a different edge set.

To this aim let be *𝓗* = (*W, F*) = *f* ^−1^(*R*_+_, *R*_*−*_). Since a rewiring does not affect the node set of the transformed network, *R*_+_ has the same node set of *B*_+_, and *R*_−_ has the same node set of *B*_−_. On the other hand, *B*_+_ and *B*_*−*_ are the two bipartite networks induced by the positive and negative edges (respectively) of *𝒢*. For construction, the union of their nodes gives *V*. Taken together these observations imply that *W* = *V*

From the definition of *f, B*_+_ contains the positive edges in *E* and *B*_*−*_ the negative edges of *E* (whose terminal nodes have been possibly relabeled). From the definition of rewiring, the edge set of *R*_+_ contains the same number of edges of *B*_+_ but at least one edge not contained in *B*_+_. Similarly the edge set of *R*_*−*_ contains the same number of edges of *B*_*−*_ and at least one edge not contained in *B*_−_. Therefore, from the definition of *f*^−1^, |*F*| = |*E*| and *F* contains at least two edges that are not included in *E*. This imply that *F* ≠ *E*.

As a conclusion *𝒢* and *𝓗* have the same set of nodes and number of edges but different edge sets. Secondly we need to show that the signed degrees of all the nodes of *𝓗* are equal to those of all the nodes in *𝒢*.

Let us assume that the positive-in-degrees of *𝓗* are different from those of *𝒢*. From the *f*^−1^ definition, this implies that *R*_+_ contains at least a node in the source set for which the degree is different from that of its counterpart in *B*_+_. However, this contradicts *R*_+_ being a rewired version of *B*_+_. With the same argument it is possible to prove that all the signed-directed node degrees of *𝓗* are equal to those of *𝒢*.

### 2.3 Switching-algorithm lower bound for bipartite networks of any size

To rewire a bipartite network *B* = (*S, D, E*), the switching-algorithm (SA) [21] performs a cascade of switching-steps (SS). In each of these SS two edges (*a, b*) and (*c, d*) are randomly selected from *E* and replaced with (*a, d*) and (*c, b*) if these two new edges are not already contained in *E*. In this case the SS under consideration is said *successful.*

Underlying the SA is a Markov chain whose states are different rewired versions of the initial network *𝒢* and a transition between states is realized by a successful SS.

In [20] we prove that, if executing a sufficiently large number of SS, the SA can efficiently simulate samplings from the uniform distribution of all the possible bipartite networks with predefined node sets and prescribed node degrees. Therefore it can be used to obtain a rewired version of a network *B* that it is, on average, no more similar to *B* than are similar to each other two bipartite networks *B*_1_ and *B*_2_ sampled from the real uniform distribution of all the possible bipartite networks with the same node sets and node degrees of *B*. To this aim, the number of SS to be performed before sampling the (*k* + 1)-th rewired network must be large enough to assure that the algorithm has *forgotten* the *k*-th sampled rewired network (the starting network *𝒢* for *k* = 0). Formally, the number of SS between two following samplings must be at least equal to the burn-in time of the Markov chain underlying the SA, which is needed to reach a stationary distribution [25, 26]. An example of this is shown in Figure 2: the 5 plots show results from a simulation study in which the SA has been used to rewire a synthetic bipartite network of 50 + 50 nodes and an edge density of 20%, and rewired versions of this network have been sampled at different intervals of SSs. A sampling interval of 1 SS produces sampled networks that are strongly related to each other (Figure 2A). Gradually increasing the sampling interval (from 5 to 20 SS, Figure 2B to D), reduces the sampled network similarities but some local dependencies are maintained. At a sampling interval of 300 SS (Figure 2E) the Markov chain underlying the SS has reached its stationary distribution, the sampled networks are completely unrelated and there are no dependencies. Therefore, for the bipartite network under consideration, a number of SS ≥300 is sufficient to simulate samplings from the uniform distribution of all the possible bipartite networks with 50 + 50 nodes and node degrees equal to those of the original network.

**Figure 2.**
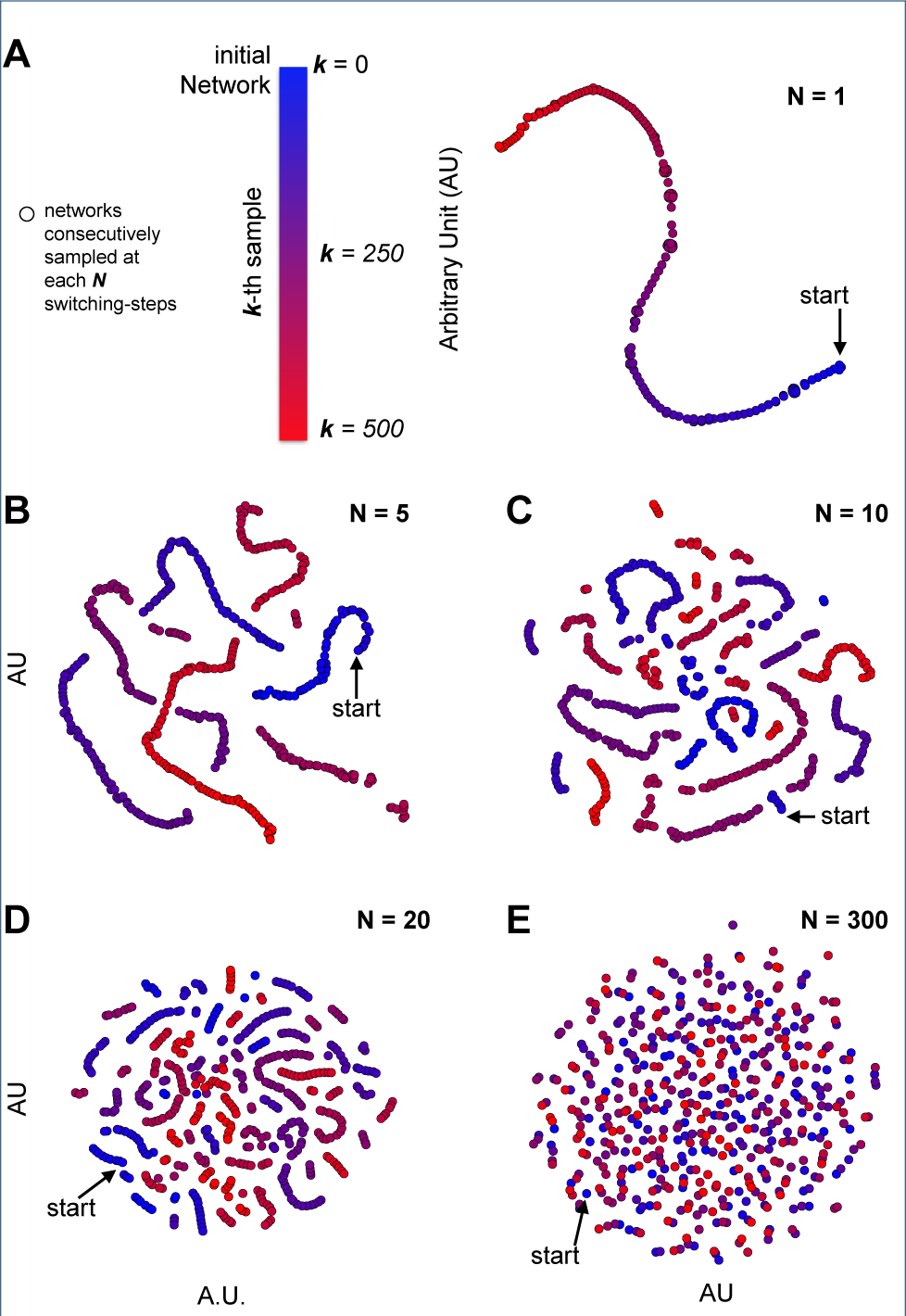
Rewired network samplings using the switching-algorithm (SA) at different sampling intervals, in terms of switching-steps (SS). Points represent sampled networks, arrows indicate a starting synthetic network, and colors indicate the sampling order. Point proximities reflect corresponding network similarities quantified through the Jaccard index. Point coordinates have been obtained with a generalized multi-dimensional scaling procedure (t-SNE).

An empirical bound *N*′ for the minimal number of SS to be performed by the SA between two consecutive samplings has been proposed in [21] as being equal to 100 times the number of edges of the bipartite network to rewire. This makes rewiring moderately large networks computationally very expensive.

By analyzing the trend of similarity to the original network along the sample path of the Markov chain simulation implemented by the SA, in [20] we proposed a novel lower bound to the number of SS needed to rewire large bipartite networks equal to

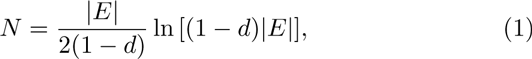

where *E* is the set of edges of the network to rewire *B* = (*S, D, E*) and d = |*E*|/(|*S*∥*D*|) is its edge density. In [20] we show that this bound is much lower than *N*^′^ and that our SA implementation and bound provide a massive reduction of the computational time required to rewire large bipartite networks (with thousands of nodes and tens of thousands of edges) with respect to other SA implementations [22] and the bound *N*^′^.

Here we provide a generalization of the lower bound *N* making the SA effective and computationally efficient in rewiring bipartite networks of any size. This is led by the observation that a DSN modeling a pathway (and the two bipartite networks induced by its positive and negative edges, respectively) can be even composed by a modest number of nodes and edges.

As shown in the supplementary data of [20] (from now on going, equations from this paper will have GSD, for Gobbi supplementary data, as prefix), Equation 1 follows from the GSD-Equation 11 (page 20) and it is a simplified form of

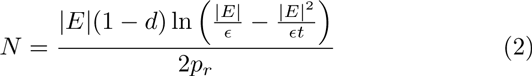

where *t* = *|S║D*| is the total number of possible edges of the original network, *d* = |*E*|/*t* is its edge density, *p_r_* is the probability of a SS to be successful. e is the accuracy of the bound in terms of distance (quantified through the convergence metric that we used to monitor the Markov chain underlying the SA, based on the number of edge shared by the original network and its rewired version at the generic *k*-th SS, and defined in GSD-Equation 9, page 19) from the real fixed point 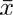.

Under the assumption of a uniform degree distribution^1^ we showed that *p_r_* = (1 − *d*)^2^ (GDS-Equation 4, page 16). As a consequence Equation 2 can be rewritten as:

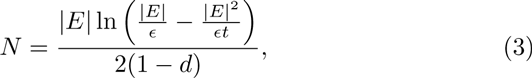

which for ∈ = 1, gives Equation 1.

Equation 3 expresses the lower bound of the number of SS as a function that accounts for the network topology and the estimated distance of the Markov chain underlying the SA from its steady-state, according to the convergence metric used in [20]. More detailed, this distance is equal to 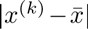, where *x*^(*k*)^ is the number of common edges between the original network and its rewired version after *k* SS, and 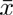 is the expected number of common edges between the original network and its rewired version, after the Markov chain underlying the SA has reached its stationary distribution.

In our previous bound definition ∈ was defined in terms of number of edges, and *N* defined as in Equation 1 in order to have 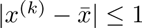 for *k* ≥*N*.

For large bipartite networks, i.e. |*E*| > 10000, a value of ∈ = 1 guarantees a relative error *δ* < 0.01% of edges for a number of SS *k* ≥ *N*. However, for relatively smaller networks, for example when |*E*| = 100, a value of ∈ = 1 implies a substantially increase in the relative error to *δ* = 1%, making the estimated lower bound *N* increasingly suboptimal with respect to the estimated real fixed point.

For this reason here we redefine the lower bound *N* for the number of SS as a function of its relative error *δ*, which quantifies its sub-optimality with respect to the estimated real fixed point. Through the simple substitution ∈ = |*E*| *δ*, Equation 3 can be rewritten as:

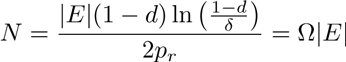

where 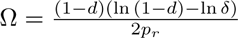 depends only on the level of accuracy *δ*, the density d of the original network and the probability *p_r_* of a successful SS. For uniformly distributed degrees^[1]^, i.e. *pr* = (1 - *d*)^2^, this bound reads as:

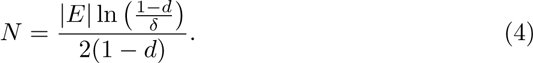

A value of *δ* = 0.00005 (corresponding to ∈ =1 edge when |*E*| ~ 20000), is used by default by our new implementation of the SA in the new version of the package *BiRewire* but this parameter can also be set to a user defined value, making our tool and bound suitable for the rewiring of bipartite networks of any size. Additionally, the choice of a suitable value for this parameter can be determined by visually inspecting the SA Markov chain convergence with a new dedicated function (described in Section 3.1)

### 2.4 Convergence criteria for signed directed networks

In [20] we showed that the convergence criteria we used to estimate our lower bound *N* for the number of switching-steps (SS) needed to rewire bipartite networks can be applied also to the more generic case of undirected networks.

To show the validity of this criteria for F-rewiring of directed signed networks (DSNs) let us observe that the Jaccard Index (*J*)[27] used to assess the similarity between two DSN with the same set of nodes and same number of edges: *𝒢* = (*V, E*) and *𝓗* = (*V, F*) is defined as

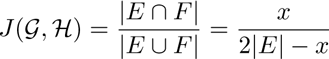

where *x* = | *E* ∩ *F*| is the number of common edges and the last equivalence holds because the two DSNs have the same number of edges. When estimated for bipartite networks, our *N* guarantees that the number of common edges between an initial network B and its rewired version at the *N*-switching-step is asymptotically minimized.

#### Proposition 2.2.

*Let be R*_+_ *and R*_-_ the *rewired versions of two bipartite networks B*_+_ *and B*_−_ *obtained through a number of switching-steps respectively equal to N*_+_ *and N*_−_ (*both computed using* Equation 4), *and such that* (*B*_+_,*B*_−_) = *f*(*𝒢*) *(where* f *is the transformation function defined in section 2.2 and* 𝒢 *a DSN). Then the Jaccard similarity between* 𝒢 *and H* = *f*^−1^(*R*_+_, *R*_−_) *is minimized.*

**Proof.** *J* (*𝒢, 𝓗*) reaches a minimum when the number of common edges *x* between *𝒢* and *𝓗* reaches a minimum. *x* is given by the sum of the number of common positive and negative edges across the two networks, namely *x* = *x*_+_ + *x*_−_. Given that *𝓗* = *f* ^−1^(*R*_+_,*R*_-_), *x*_+_ is the number of common edges between *B*_+_ and *R*_+_. Analogously *x*_-_ is the number of common edges between *B*_−_ and *R*_−_. Since *R*_+_ and *R*_-_ are rewired version of *B*_+_ and *B*_-_ computed through *N*_+_ and *N*_-_ (minimizing *x*_+_ and *x*_-_, respectively) also *x* = *x*_+_ + *x*_-_ is minimized.

## 3 Results

### 3.1 Overview of the new functions included in *BiRewire* v3.0.0

The R-package *BiRewire* **(http://bioconductor.org/packages/BiRewire/)** was originally designed to efficiently rewire large bipartite networks ([20]). We have performed a major update, by including functions to:a

- read/write directed signed networks (DSN) from/to simple interaction format (SIF) files (functions **birewire.load.dsg** and **birewire.save.dsg**);
- perform the transformation *f* (and its inverse *f*^−1^) to derive bipartite networks induced by positive and negative edges of a DSN, and viceversa (functions **birewire.induced.bipartite** and **birewire.build.dsg**);
- F-rewire a DSN by applying the switching-algorithm (SA) to the two corresponding induced bipartite networks with numbers of switching-steps automatically determined for both networks individually, using Equation 3 (function **birewire.rewire.dsg**);
- sample *K* rewired versions of a network: this function runs *K* instances of the SA in cascade; each of these instances performs a number of switching-steps (SS) determined using Equation 3. This function can take in input a bipartite network, an undirected network or a DSN (in this case Equation 3 is used individually for the two bipartite networks induced by the positive and negative edges of the DSN, respectively) (**birewire.sampler.\*** functions);
- monitor the convergence of the Markov chain underlying the SA on user defined networks. This routine samples a user-defined number of networks at user defined intervals of SS. For each of these intervals, it computes a Jaccard similarity [28] pair-wisely comparing the sampled networks to each other; finally it plots the sampled networks in a plane where points proximities reflect Jaccard similarities of the corresponding networks and point coordinates are computed through the generalized multidimensional scaling method *t-SNE* [29]; this function gives in output the network coordinates of such scale reductions and produce the plots shown in Figure 2. Also in this case the inputted graph can be a bipartite network, an undirected network or a DSN (**birewire.visual.monitoring.\*** functions);
- perform an analysis of the trends of Jaccard similarity across SS. This function performs a user-defined number of independent runs of the SA, computing the mean value and a confidence intervals of the observed pairwise Jaccard similarities between the obtained rewired networks. The result is a dataset containing the Jaccard similarity scores computed and sampled at user-defined intervals of SS, and a plot similar to that showed in Figure. 3A and 4A. This function takes in input a bipartite network or an undirected network or a DSN (**birewire.analysis.\*** functions).

**Figure 3.**
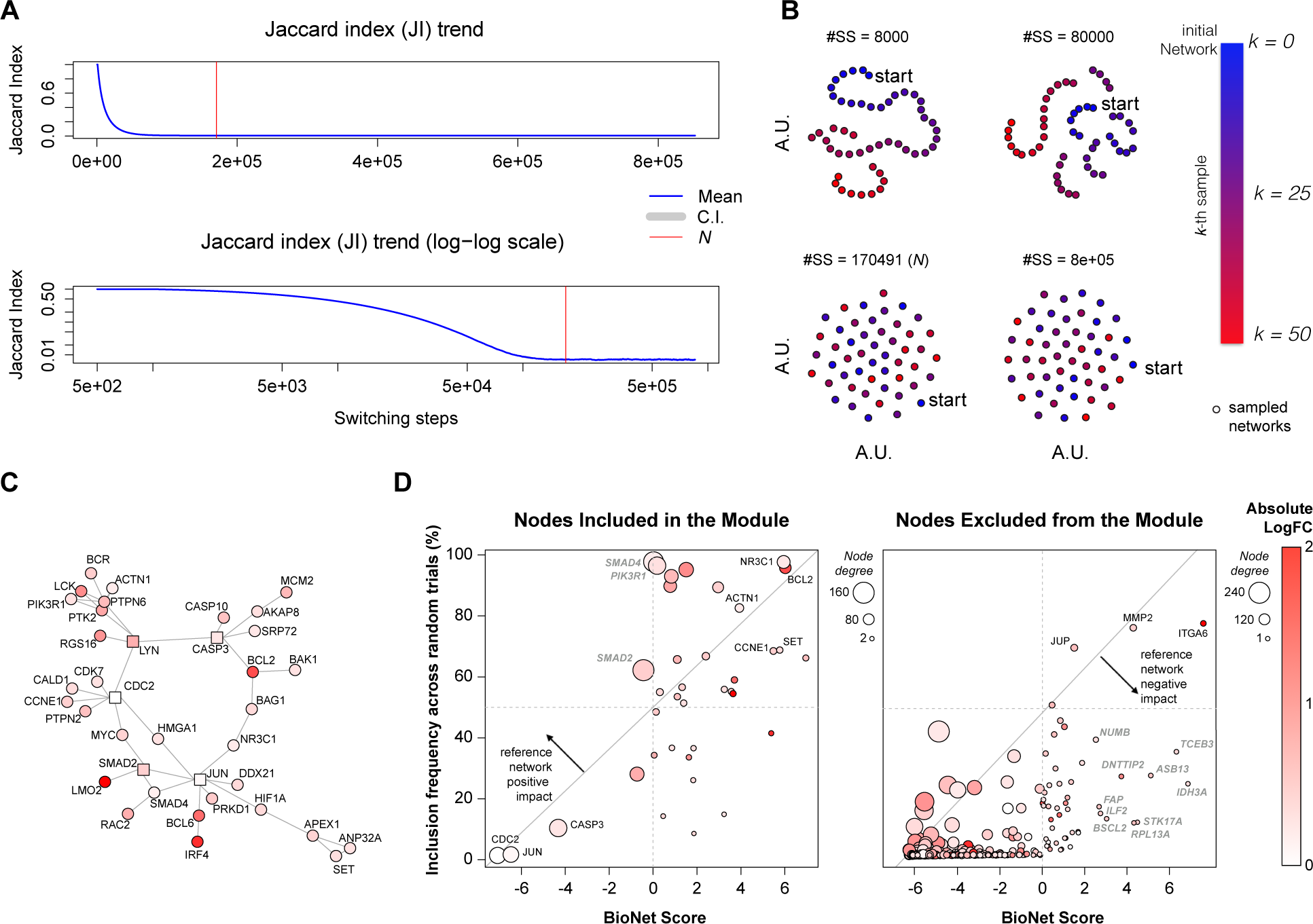
BioNet study case. (A) Analysis of the Jaccard index trend across switching-steps (SS) while rewiring the BioNet reference Interactome and estimation of the lower bound N; (B) visual inspection of the switching-algorithm Markov chain convergence to verify the suitability of the estimated bound (see Figure 2 legend for further details); (C) Interactome module outputted by BioNet while analyzing the DLBCL dataset; (D) scatter plots of BioNet scores vs. frequency of inclusion in the rewired solutions for all the nodes included in the BioNet module (left plot) and for all the other Interactome nodes contained in the DLBCL dataset (right plot).

**Figure 4.**
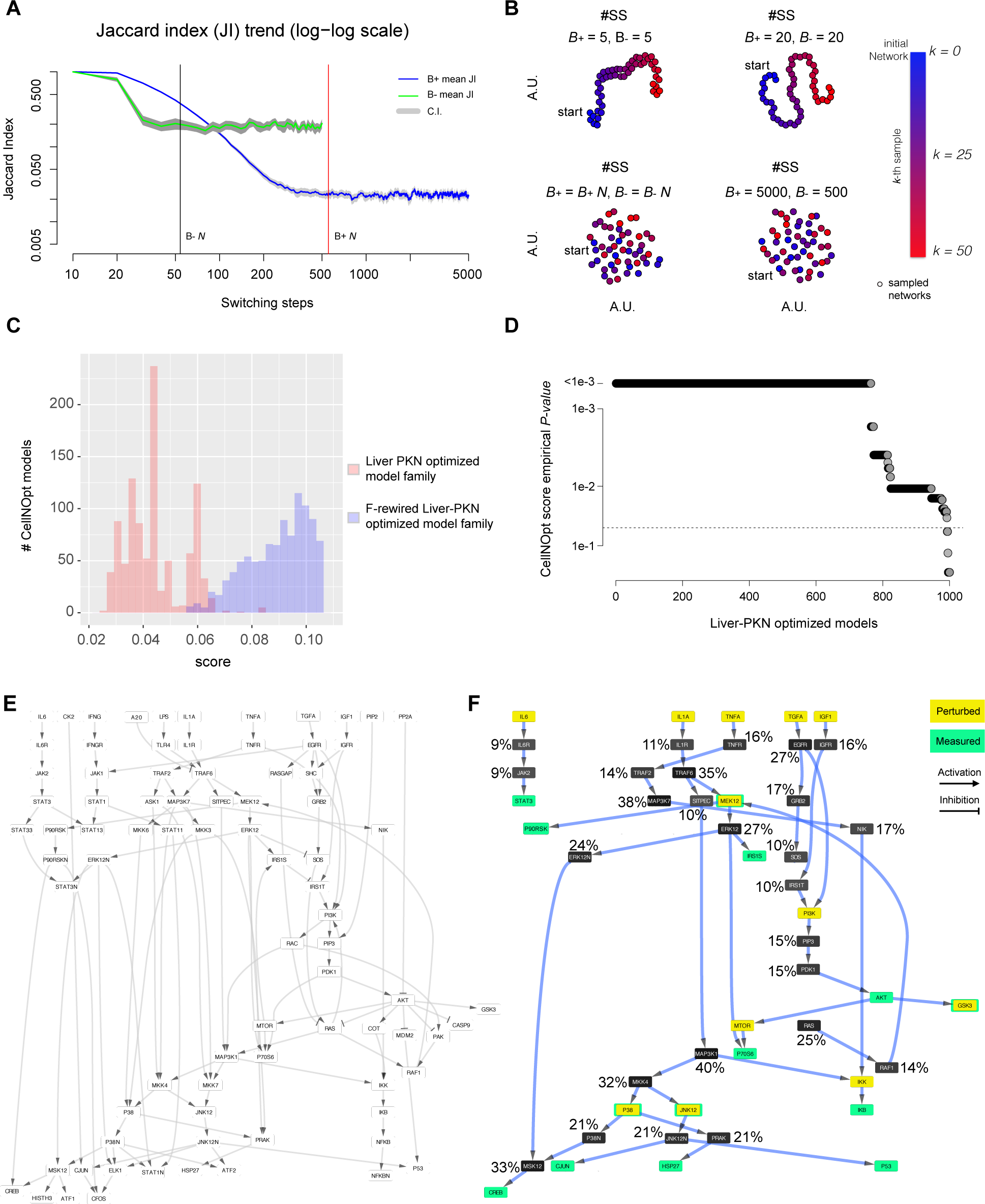
CellNOpt study case. (A) Analysis of the Jaccard index trend across switching-steps (SS) while rewiring the two bipartite network induced by the positive (respectively negative) edges of the reference DSN (liver prior knowledge network (liver-PKN)) and estimation of the lower bounds for the number of switching-steps; (B) visual inspection of the switching-algorithm Markov chain convergence to verify the suitability of the estimated bounds (see Figure 2 legend for further details); (C) Comparison of the CellNOpt scores and the rewired scores; (D) Empirical p-values of the CellNOpt scores across the entire family of models. (E) The liver-PKN used by CellNOpt as initial reference network; (F) The model outputted by CellNOpt when using the liver-PKN as initial reference network with superimposed the frequency of inclusion of each node in a set of 1,000 models outputted by CellNopt using F-rewired versions of the liver-PKN as reference networks.

Worthy of note is that, supporting the analysis of DSNs, our package can handle also generic directed graphs, therefore with *BiRewire3* it is now possible to rewire any kind of unweighted networks.

We have developed also a cython wrapper of the corresponding C library for Python users. A first release (with some basic functions) can be found in **https://github.com/andreagobbi/pyBiRewire**.

### 3.2 Case study 1: BioNet

The R package *BioNet* [30] provides a set of methods to map gene expression data onto a large reference biological network, and to identify (with a heuristic method) a maximal scoring sub-network (MSS), which a is a set of connected nodes (or *module)* with unexpectedly high levels of differential expression [31]. Several other methods moving along the same lines exist (as, among others, EnrichNet [6]). Here we focus on BioNet because it can be considered a typical example among these methods, and we show how *BiRewire3* can be used to estimate the impact of the reference network topology and the functional characterization level (FCL), i.e. sign-directed degree, of its nodes on the optimal module outputted by this tool.

The initial reference network used by BioNet (the *Interactome)* is a large undirected protein-protein-interaction network assembled from HPRD [32] and encompassing 9,392 nodes and 36,504 edges. In [30], the authors show an application of BioNet to gene expression data from a diffuse large B-cell lymphoma (DLBCL) patient dataset, with corresponding survival data. After determining gene-wise P-values for differential expression and risk-association, the authors aggregate them and fit a beta-uniform mixture model to the distribution of aggregated P-values that yields a final score (accounting for both considered factors) for each gene: the higher this score the more a gene is differentially expressed across the contrasted groups of patients. Then the methods proceeds with mapping these scores onto the Interactome nodes and, applying a heuristic method [9], it identifies a sub-network (referred to as a module) that is a sub-optimal estimate of the MSS. This module is shown in Figure 3C and the BioNet package vignette contains detailed instructions on how to reproduce this result.

To evaluate the impact of the FCLs of the Interactome nodes on the module outputted by BioNet when used on the DLBCL dataset, we generated 1,000 F-rewired versions of the Interactome with *BiRewire3* and used each of them as initial reference network in 1,000 individual BioNet runs, using the DLBCL dataset as input.

To this aim we first conducted a *BiRewire3* analysis (using the dedicated function of our package) to determine the number of switching-steps (SS) to be performed by the switching-algorithm (SA) in order to F-rewire the Interactome. This function makes use of the convergence criteria we designed in [20], which is based on the estimated time, in terms of SS, in which the Jaccard similarity (JS) between the original network and its rewired version at the *k*-th SS reaches a plateau (Figure 3A). In [20] we showed that this criteria is equivalent to other established methods to monitor Markov chain convergence when the states are networks. In addition its relatively simple formulation consents the analytical derivation of an estimated plateau time, i.e. our bound *N*. Neverthless, our package allows also a visual inspection of the optimality of the estimated bound *N* showing how independent are F-rewired versions of an initial network sampled at a number of user-defined SS intervals as well as every N SS (Figure 2).

These preliminary analyses resulted in a required number of SS equal to *N* = 170,491 (Figure 3A) and showed that this number of SS is actually sufficient to generate unrelated F-rewired versions of the Interactome, thus to simulate samplings from the uniform distribution of all the possible networks with the same number of nodes and FCLs of the Interactome (Figure 3B). Generating 1,000 F-rewired versions of the Interactome sampled each *N* SS required ~ 2 hours on a 4 core 2.4 Ghz computer with 8GB memory.

Running 1,000 independent instances of BioNet using each of these F-rewired Interactome as reference network and the DLBCL dataset in input resulted into 1,000 different module solutions (rewired solutions). For each of the nodes included in the original BioNet module solution (Figure 3C), we quantified the ratio of rewired solutions including them and we investigated how this quantity related to the corresponding BioNet scores (3D). As expected, we observed a significant correlation (*R* = 0.51, *p* = 0.001). In fact, as per the definition of the MSS, it is reasonable that nodes with high scores (such as, for example *NR3C1* and *BCL2)* tend to be included in the module outputted by BioNet regardless their edges and degree in the reference Interactome. Similarly, nodes with large negative scores (such as *CDC2* and *JUN)* are included in the module only because they link high scored nodes and it is obvious that they do not tend to be included in the rewired solutions, as in this case the way they are interlinked to other nodes is crucial.

Nevertheless, a number of nodes (such as, *SMAD4, SMAD2* and *PIK3R1)* have modest score but tend to be included very frequently in the rewired solutions. This hints that what leads the inclusion of such nodes in the BioNet module is their high FCL. As a confirmation of this, *SMAD4, SMAD2* and *PIK3R1* fall over the 99th percentile when sorting all the nodes in the Interactome (and included in the DLBCL) based on their FCL (which in this case corresponds to their degree). This is a proof that the reference network provides the BioNet outputted module with a *positive impact*, and that at least some nodes are included in the solution because of their high FCL.

When extending this analysis to the nodes of the Interactome (included in the DLBCL dataset) that are not present in the module outputted by BioNet we observed again an expected significant correlation (*R* = 0.51, *p <* 10^−16^), and some nodes (such as *JUP, MMP2* and *ITGA6)* with high scores frequently included in the rewired solutions (the fact that these nodes do not appear in the BioNet outputted module is due to the sub-optimality of the used heuristic). However we also observed a large number of nodes (such as *RPL13A, STK17A* and *IDH3A)* scored high but relatively infrequently included in the rewired solutions. This hints that these nodes are penalized by their low FCL in the reference Interactome, thus proving the existence of a *negative impact* provided by the reference Interactome to the BioNet outputted module, and that at least some nodes are not included in the solution because of their low FCL.

An indication of both these types of impacts, together with diagnostic plots and statistics would complement and complete the output of many valuable and widely used tools, such as BioNet.

### 3.3 Case study 2: CellNOpt

CellNOpt (www.cellnopt.org) is a tool used to train logic models of signal transduction starting from a reference directed signed network (DSN) called a prior knowledge network (PKN), describing causal interactions among signaling species (obtained typically from literature), and a set of experimental data (typically phosphorylation), obtained upon various perturbatory conditions ([23]).

CellNOpt converts the PKN into a logic model and identifies the set of interactions (logic gates) that best explain the experimental data. This is performed through a set of Bioconductor packages supporting a number of mathematical formalisms from Boolean models to ordinary differential equations.

Through a built-in genetic algorithm CellNOpt identifies a family of subnetworks from the reference DSN (from now, models) together with the value of the objective function (the *model score δ*) quantifying at what extent each model is able to explain the experimental data (the lower this value the better is the fit of the model to the data). By default, the best model with the lowest score denoted 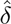 is returned to the end-users. Note, however, that multiple models may be returned if they cannot be discriminated given the experimental evidence. Besides, to account for experimental noise, users may also provide a parameter, which is called tolerance (in percentage), that will keep all models below a threshold defined as 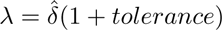.

Setting this tolerance parameter is non-trivial and depends largely on the experimental error. One idea would be to estimate this threshold by looking at the expected ability of F-rewired versions of the liver-PKN to explain the data, when they are used as input to CellNOpt. In fact, even if original local node properties are maintained, in each of these F-rewired networks the topology of the biological pathways interlinked in the liver-PKN is disrupted. As described before, a large score calculated by CellNOpt indicates a large disagreement between data and network logic behavior at the measured nodes. Therefore the distribution of the *δ*s outputted by CellNOpt when using these F-rewired networks gives an idea of the attainable *base-line performaces*, which are not derived from biologically meaningful models but depend only on the FCL (signed and directed node degrees) of the original liver-PKN.

Based on these considerations, here we show how *BiRewire3* can be used to identify such a threshold as the maximal *δ* value whose deviance from expectation is statistically significant. Similarly to the previous case study, this expectation can be empirically estimated by running a large number of independent CellNOpt runs using F-rewired versions of the initial reference signaling network and the same experimental data. Thus accounting for the effect of the node FCLs on both scores and outputted models. To this aim, we used the same reference PKN network and phospho-proteomic data used in [23], which has about 80 nodes and 120 directed and signed edges. This was a study on human liver cell and hence the network is called liver-PKN hereafter. With the *BiRewire3* package we generated (in less than 10 seconds, on a standard unix laptop) 1000 F-rewired versions of the liver-PKN, visually inspecting (as in the previous case study) the optimality of our estimated lower bound *N* for the number of switching-steps (SS) to be performed by the switching-algorithm (SA) (Figure 4AB) between one sampled F-rewired network and the following one. Subsequently we run 1000 independent instances of CellNOpt (using the CellNOptR package [23], v1.16 available on Bioconductor at **https://www.bioconductor.org/packages/CellNOptR/)** on each of these F-rewired liver-PKN networks and the same phosphoproteomic dataset (obtaining one *rewired model* per each analysis), as well as a final run using the original liver-PKN network (obtaining a family of 1000 different models). When comparing the two populations of CellNOpt scores obtained from these two analyses we observed, as expected, a notably statistically significant difference (t-test p-value < 10^−16^, Figure 4C), indicating that in the F-rewired networks the topology of the pathways originally interlinked in the liver-PKN is actually disrupted. Subsequently, using the distribution of scores of the rewired models we computed empirical p-values for the CellNOpt scores for the entire model family outputted by the final run (making use of the original liver-PKN).

For a given score *δ*_i_ corresponding to the *i*—th model of the family, an empirical p-value was set equal to the number of rewired models m such that *δm* ≥ *δ_i_* divided by 1000 (the number of tested f-rewired liver-PKNs). More than 90% of the models in the outputted family had a CellNOpt score significantly divergent from expectation (p-value < 0.05) and the estimated score threshold guaranteeing this (or a greater) divergence from expectation, thus a minimal impact of the initial liver-PKN FCLs, was equal to 0.06.

Finally, and similarly to the analysis performed in the first study case, we quantified the tendency of each of the nodes included in the final merged CellNOpt model to be included in the rewired models, finding that also in this case this is indeed proportional to the nodes’ FCL.

In summary, *BiRewire3* could be effectively used to determine a score threshold on an analytical ground, based on which meaningful models could be selected from the family outputted by CellNOpt for further analyses, and finally assemble a consensual model solution. Additionally, it could be employed to evaluate the extent of impact of the CellNOpt reference network on the topology of its outputted consensual model.

## 4 Discussion

*BiRewire3* is a one-stop tool to rewire in a meaningful way any type of unweighted networks (undirected, directed, and signed) currently used to model different datasets and relations in computational biology (including presence-absence matrices, genomics datasets, pathways and signaling networks) in an computationally efficient way. It represents a significant and formally demonstrated advance with respect to its previous version [20], whose applicability was restricted to presence/absence matrices and undirected bipartite networks. We have previously shown that, thanks to an analytically derived lower bound to the number of steps of its underlying algorithm, the computational time requirements of *BiRewire3* are vastly lower than those of other similar tools, reducing from months to minutes (on a typical desktop computer) when rewiring networks with tens of thousands of nodes and edge density ranging up to 20%. Additionally, the core algorithm underlying *BiRewire3* is based on a Markov chain procedure that could be easily parallelized in future implementations, to exploit the power of modern multi-core computer architectures, thus reducing these time requirements even further.

Our package is available as free open source software on Bioconductor and, as we showed in our case studies, it can be easily combined into computational pipelines together with a wide range of existing bioinformatics tools aiming at integrating signaling networks with experimental data.

## 5 Conclusion

We have presented a computational framework implemented in a R package that could complement existing network based tools. This will be useful for computing a wide range of constrained null models testing the significance of the solutions of these tools, and to investigate how the topology of the used reference networks can potentially bias these results.

Moreover, the range of applicability of *BiRewire3* goes beyond computational biology, and includes all those fields making use of tools from network theory, from operative research, to microeconomy, and ecological research (an example of the application of *BiRewire* application in a micro-economy and technology patent study can be found at http://arxiv.org/abs/1509.07285).

## Declarations

Ethics and consent to partecipate

Not applicable

### Consent to publish

Not applicable

### Authors’ contributions

FI, TC, MBF and JSR conceived the study and designed the implemented algorithms, AG designed all the mathematical proofs, implemented the algorithms and visualisation routines. FI wrote the manuscript. JSR supervised the study. GJ participated to the supervision of the study. All the authors revised the manuscript.

### Availability of data and materials

All the code and data objects mentioned in this manuscript and used to produce the presented results are available on Bioconductor at: https://www.bioconductor.org/packages/BiRewire/.

### List of abbreviations

- F-rewiring: network rewiring preserving the functional characterization of the individual nodes;
- DSN: Directed Signed Network;
- FCL: Functional Characterization Level;
- SA: Switching Algorithm;
- SS: Switching Steps;
- GSD: Supplementary Data of [20]
- SIF: Simple Interaction Format
- MSS: Maximal Scoring Subnetwork
- DLBCL: Diffuse Large B Cell Lymphoma
- PKN: Prior Knowledge Network

## Acknowledgements

FI has been partially supported by the European Bionformatics Institute and Wellcome Trust Sanger Institute post-doctoral (ESPOD) program.

## Footnotes

Our proof applies also to non uniform degree distributions, leading to the same conclusions for the case of directed signed networks. Here we use the uniform case for simplicity.

